# Enhanced Cholesterol-dependent Hemifusion by Internal Fusion Peptide 1 of SARS Coronavirus-2 Compared to its N-terminal Counterpart

**DOI:** 10.1101/2021.01.14.426613

**Authors:** Gourab Prasad Pattnaik, Surajit Bhattacharjya, Hirak Chakraborty

## Abstract

Membrane fusion is an important step for the entry of the lipid-sheathed viruses into the host cells. The fusion process is being carried out by fusion proteins present in the viral envelope. The class I viruses contains a 20-25 amino acid sequence at its N-terminal of the fusion domain, which is instrumental in fusion, and is termed as ‘fusion peptide’. However, Severe Acute Respiratory Syndrome Coronavirus (SARS) coronaviruses contain more than one fusion peptide sequences. We have shown that the internal fusion peptide 1 (IFP1) of SARS-CoV is far more efficient than its N-terminal counterpart (FP) to induce hemifusion between small unilamellar vesicles. Moreover, the ability of IFP1 to induce hemifusion formation increases dramatically with growing cholesterol content in the membrane. Interestingly, IFP1 is capable of inducing hemifusion, but fails to open pore.

---

Membrane fusion is a crucial step for successful entry and infection of the enveloped viruses, leading to the transfer of viral genetic materials into the host cell.^1–5^ The fusion event is triggered by the viral fusion protein that comes into action after the receptor binding domain interacts with the cell surface receptor proteins.^6^ Generally, for class I viruses, a 20-25 amino acid stretch present in the N-terminus of the fusion protein is known as fusion peptide, which is instrumental in binding with the host cell and initiating the fusion process.^7–8^ Severe acute respiratory syndrome (SARS) is an emerging form of pneumonia caused by SARS-CoVs, and the entire world is now going through a crisis due to the attack of SARS-CoV-2. The fusion domain of SARS-CoV spike protein (S2) contains three putative fusion peptides recognized as N-terminal fusion peptide (FP), internal fusion peptide 1 (IFP1), and internal fusion peptide 2 (IFP2).^9–13^ The S2 protein contains heptad repeats, HR1 and HR2, and a transmembrane region at the C-terminus in addition to these fusion peptides. Interestingly, the FP and IFP1 are highly homologous between SARS-CoV-1 and SARS-CoV-2 (**Table 1**). Therefore, proper understanding of the role of FP and IFP1 in inducing membrane fusion would provide valuable mechanistic insights of the entry of both SARS-CoV-1 and SARS-CoV-2. The atomic resolution structure of the complex formed by two heptad regions revealed the formation of a six-helix bundle; considered to facilitate close apposition of two fusing membranes.^14–15^ Membrane composition plays a significant role in the fusion process as it alters the fusion protein or peptide conformation as well as the membrane organization and dynamics.^16^ The role of cholesterol in membrane fusion is firmly established from the results obtained from viral and model membrane fusion.^17–18^ Cholesterol is also known to promote oligomerization of the SARS-CoV FP.^19^

**Table 1:** Sequences of FP and IFP1 for SARS-CoV-1, SARS-CoV-2 and peptides used in the study

| Fusion Peptide |  |
| --- | --- |
| SARS-CoV-1 | MYKTPTLKDFGGFNFSQIL |
| SARS-CoV-2 | IYKTPTLKDFGGFNFSQIL |
| Internal Fusion Peptide 1 |  |
| SARS-CoV-1 | GAALQIPFAMQMAYRF |
| SARS-CoV-2 | GAALQIPFAMQMAYRF |

The lipid stalk hypothesis assumes that the sequential evolution of the intermediates toward the opening of fusion pore. Initially, two bilayers come close and the outer leaflets of both bilayers mix to form the stalk intermediate. Subsequently, the inner leaflets of the apposed membranes come in contact with each other to form transmembrane contact, which finally undergoes mixing of inner leaflets to open fusion pore. The stalk and transmembrane contact structures are collectively called hemifusion intermediates. A schematic representation of the fusion process is shown in **scheme 1**.

**Scheme 1.**
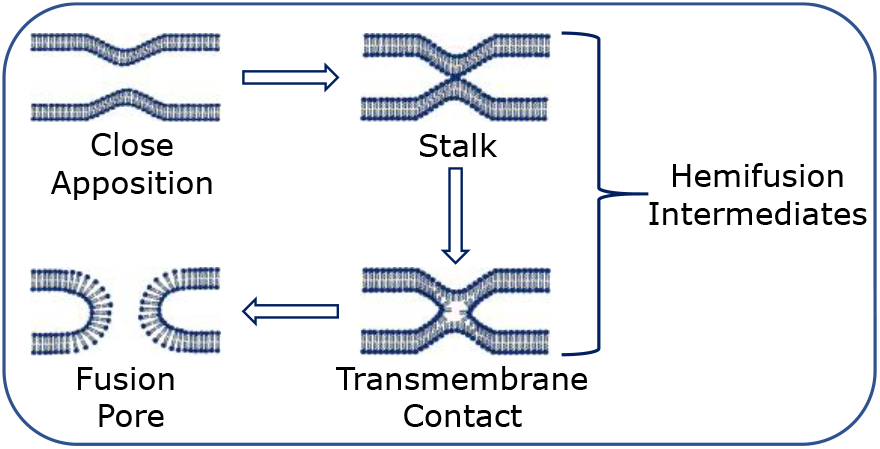
Schematic representation of different intermediates during the course of membrane fusion.

In this work, we have studied the effectiveness of FP and IFP1-induced fusion of small unilamellar vesicles (SUVs), and evaluated the effect of membrane cholesterol on the fusion process. Our results demonstrate that the IFP1 promotes lipid mixing in a cholesterol-dependent fashion. Both the rate and extent of lipid mixing increase significantly in presence of cholesterol. On the contrary, the FP is not that efficient to induce lipid mixing, however, there is a slight increase in rate and extent of lipid mixing in presence of membrane cholesterol. Interestingly, both FP and IFP1 fail to demonstrate substantial content mixing highlighting the role of other domains of S2 protein for the pore formation. The extent of content leakage remains about 10%, which confirms the overall integrity of fusing membranes.

The above observation indicates that the IFP1 (and partially FP) induces hemifusion, however incapable of opening the pore between two fusing membranes. Our results support the requirement of interaction between FP and transmembrane domain of fusion protein for pore opening as proposed earlier in HIV.^20^

In order to evaluate the effect of FP and IFP in membrane fusion, we have measured lipid mixing, content mixing, and content leakage kinetics using fluorescence-based methodologies described in method section (Supplementary material). IFP1 induced about 51% of lipid mixing in DOPC/DPOE/DOPG (60/30/10 mol%) SUVs in a lipid to peptide 100:1. The rate and extent of lipid mixing increases with increasing cholesterol concentration, and extents are about 71% and 84% in DOPC/DOPE/DOPG/CH (50/30/10/10 mol%) and DOPC/DOPE/DOPG/CH (40/30/10/20 mol%) SUVs, respectively (Figure 1A, Table-2). This result suggests that the efficiency of IFP1 in promoting lipid mixing is extremely dependent on the concentration of membrane cholesterol. Though it promotes significant amount of lipid mixing, does not induce content mixing, and brings about 10% content leakage in the membrane containing 20 mol% of cholesterol (Figure 1B&C). Putting this observation in the context of membrane fusion it is clear that the IFP1 is capable of inducing hemifusion intermediate but unable to open the fusion pore. The hemifusion is solely dependent on lipid mixing, where the lipids of outer leaflets of two fusing membranes mix with each other. A small amount of content mixing in the hemifusion intermediate is possible as the small fluorophores can move from one membrane to the other through the thermal fluctuation. Moderately low content leakage indicates the overall integrity of the membrane during the formation of hemifusion intermediates. Interestingly, the content leakage data saturates within about 400 seconds, which designates that the content leakage is majorly observed during the lipid reorganization forming the hemifusion intermediate. Similar experiments were carried out in three different lipid compositions with the N-terminal FP, and the results are shown in Figure 2(A-C). The FP promotes nominal amount of lipid mixing in all three lipid compositions in a lipid to peptide ratio of 100:1. The extent of content mixing and content leakage are similar to what we observed in presence of IFP1. Overall, our result suggests that the N-terminal FP is less efficient in promoting hemifusion, FP does not rupture the membrane as evident from the moderately low content leakage, and content leakage majorly takes place during the formation of the hemifusion intermediate.

**Table 2:** The extent and rate constant of lipid mixing in presence of FP and IFP1 in different lipid compositions.

| Lipid Composition | Peptide | Lipid Mixing (%) | k (Sec<sup>-1</sup>) |
| --- | --- | --- | --- |
| DOPC/DOPE/DOPG (60/30/10) | IFP1 | 50.8 | $1.3 \times 10^{-3}$ |
| | FP | 3.5 | $8.8 \times 10^{-5}$ |
| DOPC/DOPE/DOPG/CH (50/30/10/10) | IFP1 | 71.4 | $2.0 \times 10^{-3}$ |
| | FP | 8.9 | $6.5 \times 10^{-4}$ |
| DOPC/DOPE/DOPG/CH (40/30/10/20) | IFP1 | 83.6 | $2.3 \times 10^{-3}$ |
| | FP | 11.5 | $8.2 \times 10^{-4}$ |

**Figure 1.**
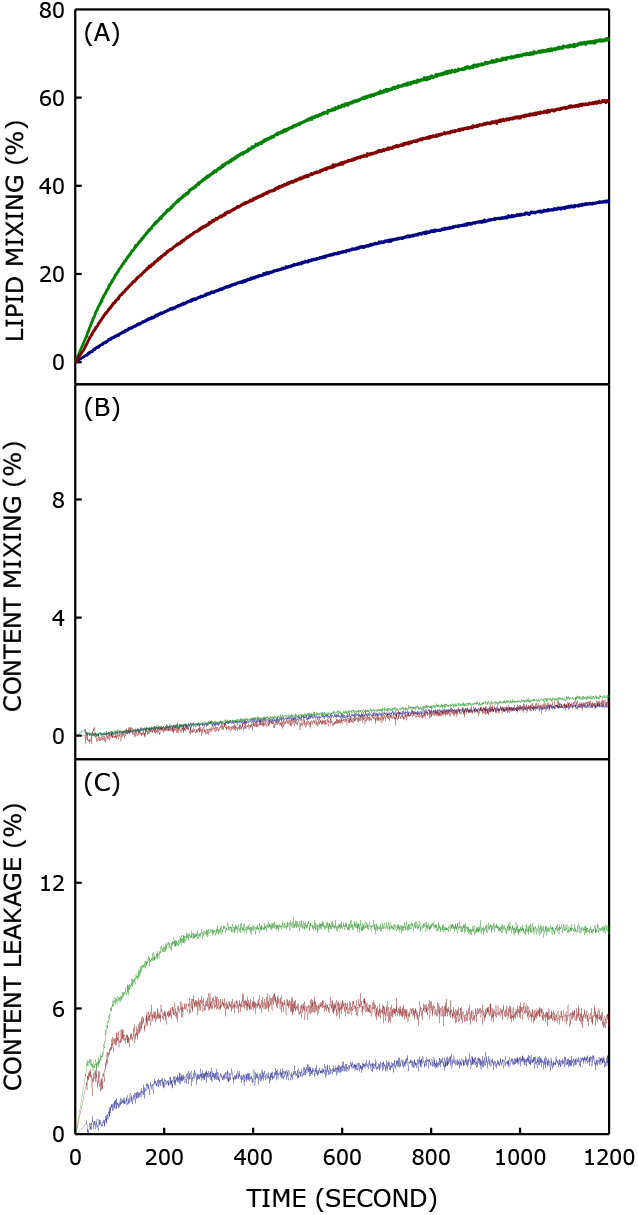
Effect of **SARS-CoV IFP1** on the kinetics of (A) lipid mixing, (B) content mixing and (C) content leakage in SUVs containing 0 mol% (Blue), 10 mol% (Red) and 20 mol% (Green) of cholesterol at 37 °C, keeping lipid to peptide ratio of 100:1. See the Materials and methods section for more details.

**Figure 2.**
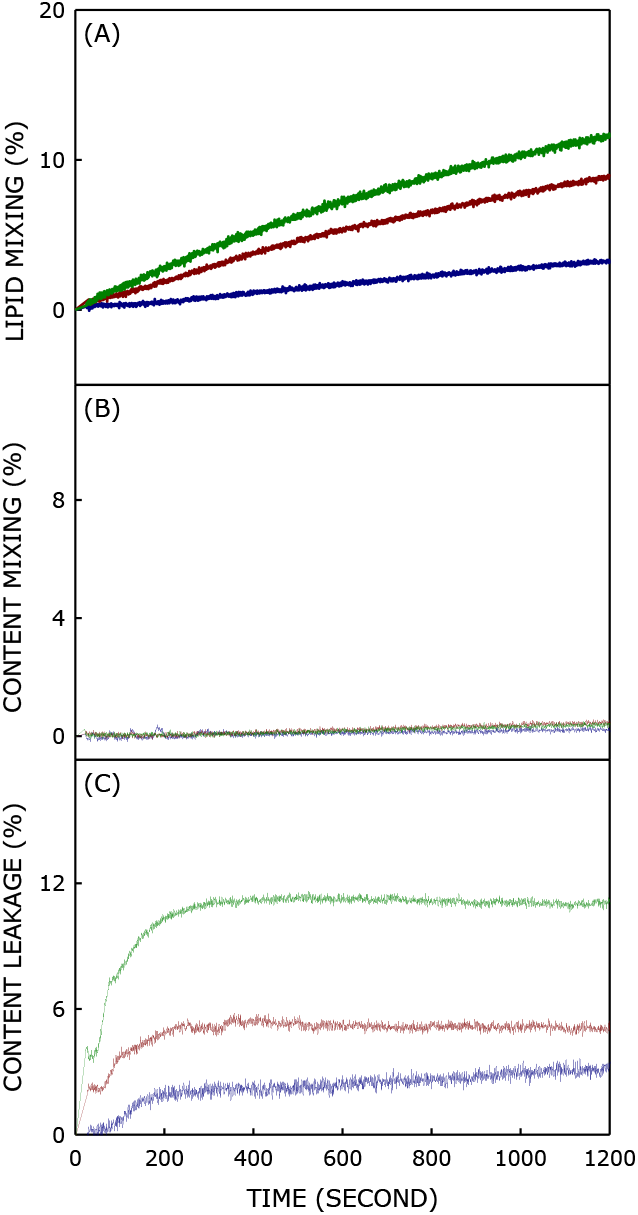
Effect of **SARS-CoV FP** on the kinetics of (A) lipid mixing, (B) content mixing and (C) content leakage in SUVs containing 0 mol% (Blue), 10 mol% (Red) and 20 mol% (Green) of cholesterol at 37 °C, keeping lipid to peptide ratio of 100:1. See the Materials and methods section for more details.

Generally, for the entry of class I viruses the N-terminal FP is considered to be crucial. Though SARS-coronaviruses belong to class I category, our results demonstrated that IFP1 is more fusogenic than its N-terminal counterpart. The higher fusogenicity of IFP1 could be correlated to its higher hydrophobicity compared to the N-terminal FP (**Figure 3**). Ten out of sixteen (62.5%) amino acids are hydrophobic in IFP1, whereas seven out of nineteen (36.8%) amino acids are hydrophobic in FP as per the Kyte-Doolittle hydrophobicity scale.^21^ Our results further demonstrated the important role of cholesterol in the enhancement of IFP1 and FP-induced hemifusion; an important link between the membrane cholesterol and higher risk of viral infection. The stringency of cholesterol in class I viral infection has been shown earlier, and our results indicate that the higher fusogenicity could be due to the higher effectiveness of fusion peptides in inducing hemifusion intermediate in presence of cholesterol. Cholesterol might promote membrane fusion either by modulating the peptide conformation^22,23^ and depth of penetration^18^ or changing membrane physical properties such as intrinsic negative curvature and stiffness.^24^ Cholesterol has an inverted cone like structure that generates intrinsic negative curvature to the membrane, which promotes the formation of nonlamellar fusion intermediates. In addition, cholesterol enhances overall membrane stiffness which provides mechanical stability to the highly curved intermediate structures

**Figure 3.**
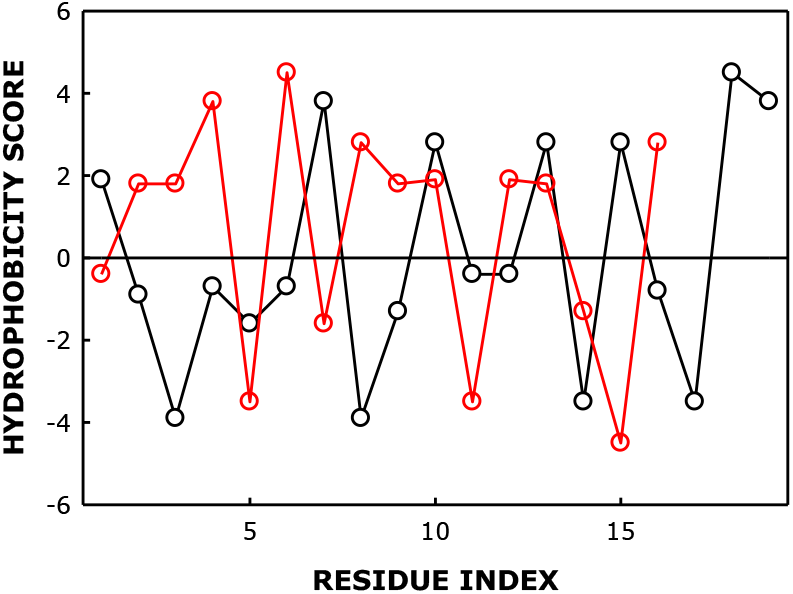
Hydrophobicity scores of each amino acid of IFP1 (Red) and FP (Black) have been plotted against the residue index. Hydrophobicity scores have been taken from Kyte-Doolittle scale.

In spite of being so successful in inducing hemifusion, both IFP1 and FP fail to open fusion pore between two fusing membranes. It was shown that the fusion peptide interacts with the transmembrane domain of the fusion protein to open up the pore.^20^ The limited ability of the fusion peptides to open up pore in our study further supports the hypothesis of interaction between fusion peptide and transmembrane domain to open the fusion pore.

Taken together, our work provides three important information regarding the fusion peptide-induced membrane fusion for SARS-coronaviruses. Firstly, it is clearly demonstrated that the IFP1 is more fusogenic than the FP and it could be due to higher hydrophobicity of IFP1. Secondly, the importance of cholesterol in the peptide-induced membrane fusion, and finally the requirement of interaction between fusion peptide and transmembrane domain for pore opening.

## ASSOCIATED CONTENT

The materials and methods are in Supporting Information. This material is available free of charge via the Internet at http://pubs.acs.org.

## AUTHOR INFORMATION

### Author Contributions

GPP: Performed all the experiments, analyzed the results and wrote the initial draft of the manuscript. SB and HC: Conceptualized the research, analyzed the results and wrote the final manuscript.

## Notes

The authors declare that there is no conflict of interest.

## ACKNOWLEDGMENT

This work was supported by research grant from the Department of Science and Technology, Government of Odisha awarded to HC. SB acknowledges support from the Ministry of Education (MOE, RG11/12), Singapore. H.C and G.P.P thanks the University Grants Commission (UGC) for UGC-Assistant Professor position, and Council for Science and Industrial Research, Government of India for Senior Research Fellowship, respectively. We acknowledge Department of Science and Technology, New Delhi and UGC for providing instrument facility to the School of Chemistry, Sambalpur University under the FIST and DRS programs, respectively. We thank Dr. S. N. Sahu of School of Chemistry, Sambalpur University, Dr. S. Haldar of Gandhi Institute of Technology and Management, and members of Chakraborty laboratory for their comments and discussions.

## ABBREVIATIONS

FP: fusion peptide
IFP1: Internal fusion peptide 1
IFP2: Internal fusion peptide 2

---

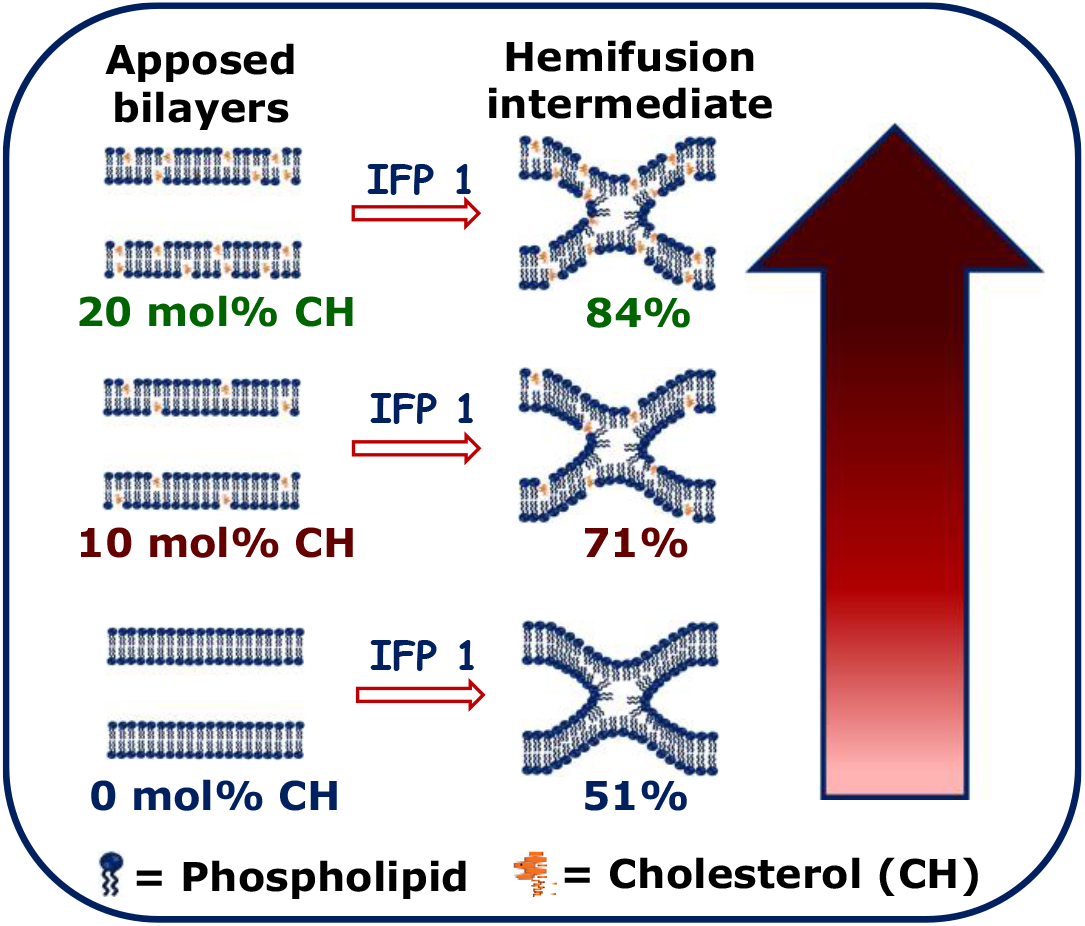

