## Supplementary material for "Enhanced Cholesterol-dependent Hemifusion by Internal Fusion Peptide 1 of SARS Coronavirus-2 Compared to its N-terminal Counterpart": Material and Methods

### MATERIALS AND METHODS

#### Materials

1,2-dioleoyl-*sn*-glycero-3-phosphocholine (DOPC),1,2-dioleoyl-*sn*-glycero-3-phosphoethanolamine (DOPE), chloroform stock solutions of 1,2-dioleoyl-*sn*-glycero-3-phospho-(1'-*rac*-glycerol) (sodium salt) (DOPG), and Cholesterol (CH) were purchased from Avanti Polar Lipids (Alabaster, AL). NBD-PE and Rh-PE were purchased from Invitrogen (Eugene, OR). Terbium chloride and N-[tris(hydroxymethyl) methyl]2-2-aminoethane sulfonic acid (TES) was procured from Alfa Aeser (Haverhill, MA). We have obtained Dipicolinic acid (DPA), Sephadex G-75, and Triton X-100 (TX-100) from Sigma Aldrich (St. Louis, MO). Calcium chloride and sodium chloride were purchased from Merck, India, and Fisher Scientific (India), respectively. Ethylenediaminetetraacetic acid (EDTA) and Spectroscopic grade DMSO were procured from Spectrochem (India). All other chemicals used in the work were of the highest available purity. Water was purified through Millipore (Bedford, MA) Milli-Q water purification system.

#### Peptide Synthesis

The fusion peptide(FP) and internal fusion peptide (IFP) of SARS-CoV was purchased commercially from GL-Biochem (China) with a purity of >98%. The peptide sequence of SARS-FP is MWKTPTLKYFGGFNFSQIL, and SARS-IFP is GAALQIPFAMQMAYRF without any modification in the N- and C-termini. Small aliquots of peptide stock solutions, prepared in DMSO, were added to the vesicle suspensions. The amount of DMSO was always less than 1% (v/v) such that it had no detectable effect on either fusion or membrane structure.

**Preparation of Vesicles**

Vesicles were prepared from either a mixture of DOPC/DOPE/DOPG (60/30/10 mol%), DOPC/DOPE/DOPG/CH (50/30/10/10 mol%), or DOPC/DOPE/DOPG/CH (40/30/10/20 mol%) using the sonication method.[1] The average hydrodynamic radii of the vesicles are 40–50 nm with a polydispersity index less than 0.2. Lipids at this appropriate molar ratio in chloroform were dried overnight in a vacuum desiccator. The dried film was hydrated and vortexed in assay buffers for 1 hour. The column (Sephadex G-75) and experimental buffer contained 10 mM TES, 100 mM NaCl, 1 mM EDTA, 1mM $CaCl_2$ at pH 7.4. Then small unilamellar vesicles were prepared using probe sonication as documented previously. All lipid mixing, contents mixing and leakage experiments were performed with 200 µM lipid. Small aliquots of peptide and probes were added from their respective stock solutions prepared in DMSO to prepare the working solutions. The DMSO content was always less than 1% (v/v), and it has been found that this small quantity of DMSO had no detectable effect on membrane structure and peptide interaction with the membrane.[2]

**Lipid Mixing Assay**

The lipid transfer during vesicle fusion was monitored using the change in FRET efficiency between NBD-PE (donor) and Rh-PE (acceptor).[3] To measure the kinetics of lipid transfer (mixing) FRET dilution was measured as a function of time as discussed earlier. In short, we prepared a set of vesicles that contain FRET pair probes in equal concentration (0.8

mol%), and hence this condition shows maximum FRET. These probe-containing vesicles were mixed with probe-free vesicles at a ratio of 1:9.[4] To evaluate the effect of SARS-FP and SARS-IFP, 2 µM peptide were added to 200 µM lipid vesicles (a mixture of probe-containing and probe-free (1:9) vesicles) to induce lipid mixing and was measured by monitoring the reduction in FRET via the change in donor fluorescence intensity. The emission intensities of the donor were monitored in Hitachi F-7000 (Japan), spectrofluorometer. The excitation and the emission wavelength for the donor (NBD-PE) are 460 nm and 530 nm, respectively. A minimum slit of 5 nm was used in both the excitation and emission side throughout the experiment. Each experiment was repeated at least three times. The percentage of lipid mixing was calculated using the following equation,[5]

$$\%\ LipidMixing = \left(\frac{F_t - F_0}{F_\infty - F_0}\right) * 100 \qquad \textbf{(1)}$$

where '$F_0$', '$F_t$', $F_\infty$' are the fluorescence intensities at the zeroth time, time = t and time = ∞, respectively. $F_\infty$ has been measured in the presence of TX-100, which is considered as the complete mixing of lipids.

**Content mixing assay**

We have monitored the content mixing as proposed by Wilschut *et al.* using $Tb^{3+}$ and DPA.[6-7] Vesicles were prepared either in 80 mM DPA or 8 mM $TbCl_3$. The untrapped DPA and $TbCl_3$ were removed from the external buffer of the vesicles using a Sephadex G-75 column equilibrated with assay buffer (10 mM TES, 100 mM NaCl, 1 mM EDTA,1 mM $CaCl_2$ at pH 7.4). 2 µM peptides were added to 200 µM lipid vesicles (mixture (1:1) of $Tb^{3+}$ and DPA-containing vesicles) to induce content mixing and study the effect of SARS-FP and SARS-IFP on content mixing. The content mixing was measured in terms of an increase in fluorescence intensity due to the formation of the Tb/DPA complex with time. The excitation and the emission wavelength for content mixing assay were used as 278 nm and 490 nm, respectively. A minimum slit width of 5 nm in both excitation and emission side was used for all measurements. Each experiment was repeated at least three times. The percentage of content mixing was calculated in the following way:

$$\% \, ContentMixing = \left(\frac{F_t - F_0}{F_\infty - F_d}\right) * 100 \qquad \textbf{(2)}$$

where '$F_0$' and '$F_t$' are the fluorescence intensities at the zeroth time and time = t, respectively. '$F_\infty$' and '$F_d$' have been calculated from the fluorescence intensity of the leakage sample at the zeroth time (which gives the highest intensity as all Tb are being complexed with DPA) and in presence of detergent, respectively. '$F_\infty$' implies the highest fluorescent signal when $Tb^{3+}$ and DPA are in the same vesicular compartment and all $Tb^{3+}$ are in complexed form with DPA. Upon detergent addition, $Tb^{3+}$ and DPA will be coming out from the vesicular compartment and being diluted in the bulk solution, resulting in dissociation of the complex and minimum fluorescence intensity ($F_d$).

**Leakage assay**

The leakage assay was carried out by monitoring a decrease in fluorescence intensity of the vesicles containing both $TbCl_3$ and DPA peptide.[6-7] 8 mM $TbCl_3$ (prepared in 10 mM TES and 100 mM NaCl, pH 7.4) and 80 mM DPA (prepared in 10 mM TES, pH 7.4) were co-encapsulated in the vesicles, and the external $TbCl_3$ and DPA were removed using Sephadex G-75 column equilibrated with assay buffer (10 mM TES, 100 mM NaCl, 1 mM EDTA, 1 mM $CaCl_2$ at pH 7.4).[1] 2 µM peptide was added to 200 µM lipid $Tb^{3+}$ and DPA containing vesicles to induce content leakage. The maximum leakage (100%) was characterized by the fluorescence intensity of a co-encapsulated $TbCl_3$/DPA vesicle treated with 0.1% (w/v) Triton X-100. The excitation and the emission wavelength were fixed at 278 nm and 490 nm, respectively. Slits of 5 nm were used in both the excitation and emission sides throughout the experiment. Each experiment was repeated at least three times. The percentage of content leakage was calculated in the following way:

$$\% \, ContentLeakage = \left(\frac{F_0 - F_t}{F_0 - F_d}\right) * 100 \qquad \textbf{(3)}$$

where '$F_0$' and '$F_t$' and '$F_d$' are the fluorescence intensities at the zeroth time, time = t, and in presence of detergent, respectively.
